# YAP is involved in replenishment of granule cell progenitors following injury to the neonatal cerebellum

**DOI:** 10.1101/558742

**Authors:** Zhaohui Yang, Alexandra L. Joyner

**Affiliations:** Biochemistry, Cell and Molecular Biology Program, Weill Cornell Graduate School of Medical Sciences, New York, NY, 10065; Developmental Biology Program, Sloan Kettering Institute, New York, NY, 10065

**Author notes:** Corresponding author: Alexandra L. Joyner, Developmental Biology Program, Sloan Kettering Institute, 1275 York Avenue, Box 511, New York, NY 10065.

**Keywords:** Hippo signaling, Nestin-expressing progenitors, granule cell precursors, Yes-associated protein, development

## Abstract

The cerebellum (CB) undergoes major rapid growth during the third trimester and early neonatal stage in humans, making it vulnerable to injuries in pre-term babies. Experiments in mice have revealed a remarkable ability of the neonatal CB to recover from injuries around birth. In particular, recovery following irradiation-induced ablation of granule cell precursors (GCPs) involves adaptive reprogramming of Nestin-expressing glial progenitors (NEPs). Sonic hedgehog signaling is required for the initial step in NEP reprogramming; however, the full spectrum of developmental signaling pathways that promote NEP-driven regeneration is not known. Since the growth regulatory Hippo pathway has been implicated in the repair of several tissue types, we tested whether Hippo signaling is involved in regeneration of the CB. Using mouse models, we found that the Hippo pathway transcriptional co-activator YAP (Yes-associated protein) but not TAZ (transcriptional coactivator with PDZ binding motif) is required in NEPs for full recovery of the CB following irradiation one day after birth. The size of the adult CB, and in particular the internal granule cell layer produced by GCPs, is significantly reduced in mutants, and the organization of Purkinje cells and Bergmann glial fibers is disrupted. Surprisingly, the initial proliferative response of *Yap* mutant NEPs to irradiation is normal and the cells migrate to the GCP niche, but then undergo increased cell death. Loss of *Yap* in NEPs or GCPs during normal development leads to only mild defects in differentiation. Moreover, loss of *Taz* does not abrogate regeneration of GCPs by *Yap* mutant NEPs or alter development of the cerebellum. Our study provides new insights into the molecular signaling underlying postnatal cerebellar development and regeneration.

## INTRODUCTION

The cerebellum (CB) not only has a principal role in motor coordination and balance control (Huang et al., 2013), but also is linked to a wide range of higher order cognitive and social functions (Fatemi et al., 2012; Marek et al., 2018; Steinlin, 2007; Stoodley et al., 2017; Tavano et al., 2007; Tsai et al., 2012; Tsai et al., 2018; Wang et al., 2014). Development of the CB in both human and mouse is a protracted process, much of which spans from late embryonic to early postnatal stages (Altman and Bayer, 1997; Dobbing and Sands, 1973; Rakic and Sidman, 1970). Therefore, the CB is particularly vulnerable to clinical and environmental insults around birth. Indeed, preterm birth has been linked to cerebellar hypoplasia and multiple neurological dysfunctions (Allen, 2008; Tam, 2013; Wang et al., 2009; Wang et al., 2014). Thus it is crucial to determine whether the CB has the capability of self-repair, and if so to dissect the molecular mechanisms that underlie such a recovery process.

The mouse CB originates from the anterior hindbrain and undergoes substantial growth in the first two postnatal weeks. Prior to birth, the *Atoh1*-expressing granule cell precursors (GCPs) migrate over the surface of the CB at embryonic day (E) 13.5–15.5. After birth, GCPs rapidly proliferate in the external granule cell layer (EGL) in response to Sonic Hedgehog (SHH) secreted by Purkinje cells (PCs) (Corrales et al., 2006; Lewis et al., 2004; Sillitoe and Joyner, 2007). Until approximately postnatal day (P) 15, post-mitotic GCPs migrate down Bergmann glial fibers and past the underlying Purkinje cell layer (PCL) to populate the internal granule cell layer (IGL) and complete differentiation and maturation (Sillitoe and Joyner, 2007). By following up on an earlier finding in infant rats that the EGL is rapidly reconstituted several days after depletion by irradiation (Altman et al., 1969), we found that the CB of neonatal mice is capable of substantial recovery and production of an adult CB with 70-80% of the normal size and normal morphology after significant ablation of GCPs in the EGL by irradiation at postnatal day (P) 1 (Wojcinski et al., 2017). Furthermore, PCs are completely replenished when ~50% are killed at P1, in an age-dependent process that involves a rare immature PC population proliferating and producing new PCs (Bayin et al., 2018). Thus, at least two cell types in the CB are effectively replenished when ablated soon after birth, and each involves a distinct cellular process.

The regeneration of GCPs after depletion via irradiation is dependent on a subpopulation of *Nestin*-expressing progenitors (NEPs), which are derived from the ventricular zone during late-embryogenesis and proliferate in the postnatal CB (Buffo and Rossi, 2013; Fleming et al., 2013; Milosevic and Goldman, 2004). There are two main subpopulations of NEPs, one that resides in PCL and gives rise to astroglia (astrocytes and Bergmann glia) and the other in the white matter (WM) produces interneurons and astrocytes (Fleming et al., 2013; Li et al., 2013; Parmigiani et al., 2015; Wojcinski et al., 2017). Furthermore, both populations rely on SHH for their proliferation. When GCPs in the EGL are depleted genetically or by irradiation at P1, NEPs in the PCL sense the injury and respond by increasing cell proliferation and then migrating into the EGL where they switch cell fate to produce GCs that contribute to the IGL (Andreotti et al., 2018; Jaeger and Jessberger, 2017; Wojcinski et al., 2017; Wojcinski et al., 2019). At the same time, NEPs in WM and IGL transiently reduce their production of interneurons and astrocytes. As expected, SHH signaling is required for NEPs to replenish the injured EGL (Wojcinski et al., 2017). The full molecular repertoire underlying the regenerative capacity of NEPs however remains to be determined.

The Hippo signaling pathway is a key regulator of size control of many organs (Halder and Johnson, 2011; Pan, 2010) through regulating cell proliferation and apoptosis (Udan et al., 2003; Wu et al., 2003). In mammals, upon activation of a conserved kinase cascade consisting of the serine/threonine kinases MST1/2 (mammalian Ste2-like kinases) and LATS1/2 (large tumor suppressor kinase 1/2), the transcriptional cofactors Yes-associated protein (YAP) and transcriptional coactivator with PDZ binding motif (TAZ, also called WWTR1, for WW-domain containing transcription regulator 1) are phosphorylated, sequestered by 14-3-3 in the cytoplasm, and targeted for degradation in a ubiquitin-proteasome-dependent manner (Callus et al., 2006; Chan et al., 2005; Wu et al., 2003; Zhao et al., 2010). Conversely, in the absence of Hippo signaling, dephosphorylated YAP and TAZ translocate into the nucleus and form complexes with the TEAD/TEF family transcription factors to activate downstream transcriptional programs that include promotion of cell proliferation and organ growth (Wang et al., 2009). Although YAP is known to regulate the renewal of many tissue types after injury, including liver, incisors, and the colonic epithelium (Hu et al., 2017; Lu et al., 2018; Yui et al., 2018), the role of YAP in mammalian brain development and regeneration remains poorly studied. Conditional genetic ablation of *Yap* in radial glial progenitor cells (using *Nestin-Cre*) was found to cause hydrocephalus and a subtle defect in the proliferation of cortical neural progenitors, but no major anatomical changes in the brain (Park et al., 2016). On the other hand, inactivation of LATS1/2 from the same (Nestin+) neural progenitor population during brain development in mouse was shown to result in YAP/TAZ-driven global hypertranscription with upregulation of many target genes related to cell growth and proliferation, as revealed by cell-number normalized transcriptome analysis (Lavado et al., 2018). The gene expression changes are thought to in turn inhibit the differentiation of neural progenitors and promote transient over-proliferation of neural progenitors. In an *ex vivo* model, it was shown that *Yap* over-expression increases the proliferation of cultured GCPs, while shRNA knockdown of *Yap* decreases proliferation (Fernandez et al., 2009). Moreover, *Yap* over-expression was reported to protect cultured GCPs from irradiation-induced damage by sustaining their proliferation and survival (Fernandez et al., 2012). However, the potential function of YAP in development and regeneration of the neonatal CB has not been tested *in vivo*.

Here we utilized genetic deletion of *Yap* and/or *Taz* in mice and revealed an essential role for YAP in mammalian CB regeneration and only a minor role in inhibiting differentiation during CB development. Loss of *Yap* at P0 from NEPs disrupted restoration of cerebellar size with pronounced reduction in the IGL and disorganization of PCs and Bergmann glial fibers. The poor recovery of the CB was associated with elevated death of the NEPs in the PCL one day after irradiation and later death of NEPs after they entered the EGL. In contrast, loss of *Yap* at P0 in NEPs or GCPs in conditional knockout (cKO) mice resulted in only mild enhancement of differentiation of NEPs and GCPs and no alteration of the size or morphology of the adult CB. Surprisingly, loss of *Taz* in addition to *Yap* at P0 in NEPs did not alter development or restrict the recovery of the CB after EGL injury. Our study identifies Hippo signaling as a key molecular signaling pathway underlying the late stage of regeneration of the postnatal CB. Our discovery also raises the possibility that inhibiting Hippo signaling could reduce cerebellar hypoplasia after injury through enhancing recovery.

## RESULTS

### YAP and TAZ expression are enriched in NEPs in the neonatal CB

As a first step in studying the function of YAP in the developing and regenerating CB, we characterized the expression of the protein in *Nestin-CFP* reporter mice using immunofluorescence (IF). It was previously reported that YAP is expressed in GCPs (Fernandez et al., 2009), but expression in NEPs was not addressed. In the CB of normal P4 mice, a low level of nuclear YAP was detected in some NEPs (CFP+ cells) whereas in GCPs in the EGL little or no YAP was detected (Fig. 1A,A’,A”). Since NEPs contribute to regeneration of the irradiated (IR) mouse CB, we asked whether YAP expression changes in NEPs during their adaptive reprogramming following irradiation of the CB. When the cerebella of mice were irradiated at P1 (IR mice) and analyzed at P4, nuclear YAP was detected mainly in NEPs (CFP+ cells) and little expression was seen in GCPs (Fig. 1B,B’,B”). Intriguingly, in IR mice most of the CFP+ NEPs that had entered the EGL showed nuclear expression of YAP (Fig. 1B,B’,B”), indicating that YAP could play a role in the adaptive reprogramming response of NEPs to EGL ablation. Unlike YAP, nuclear located TAZ expression was obvious in NEPs of both Non-IR and IR cerebella, in addition to in the NEPs that had entered the EGL (Fig. 1C-D,C’-D’,C”-D”).

**Figure 1.**
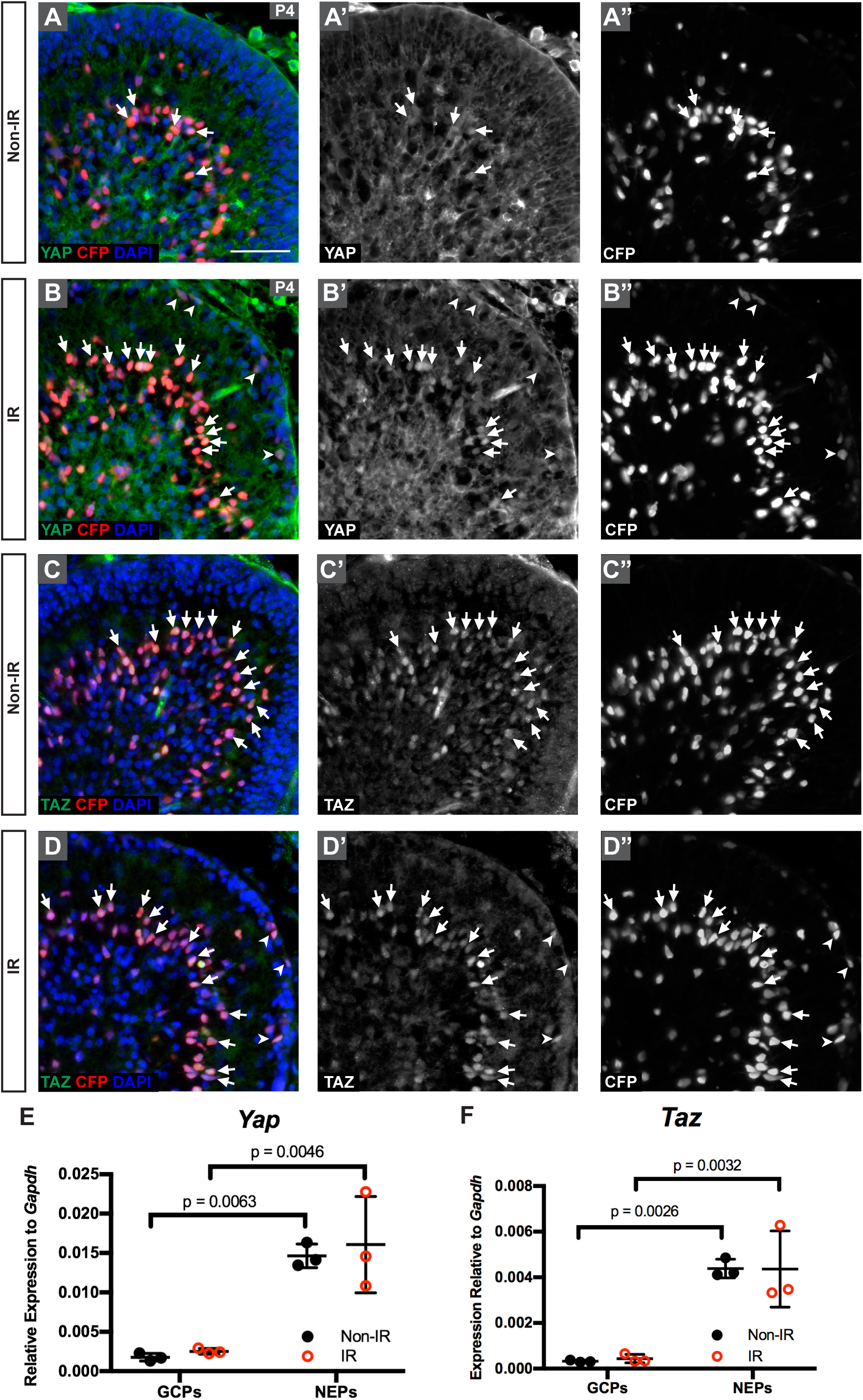
Nuclear YAP is mainly detected in NEPs in the postnatal CB. (A-D) Immunofluorescence (IF) detection of YAP (A-B) and TAZ (C-D), co-stained for CFP and DAPI, on midsagittal sections of cerebella from Non-IR and IR *Nestin-CFP* reporter mice at P4. Arrows and arrowheads indicate *Nestin* (CFP) positive cells in the Purkinje cell layer (PCL) or EGL, respectively that also have nuclear YAP or TAZ. Scale bar, 50 mm. (E-F) qRT-PCR analysis of the mRNA expression of *Yap* (E) and *Taz* (F) relative to *Gapdh* in FACS isolated NEPs (Nestin-CFP+) and GCPs (Atoh1-GFP+) from Non-IR and IR mice at P4. Data are presented as mean ± S.D., and statistical analysis by two-way ANOVA. Each data point represents one animal.

To confirm that *Yap* and *Taz* are expressed at higher levels in NEPs than GCPs, we carried out quantitative RT-PCR (qRT-PCR) of mRNA from FACS-sorted Nestin-CFP+ NEPs and Atoh1-GFP+ GCPs from P4 IR and Non-IR mice. Indeed, *Yap* and *Taz* mRNA were significantly lower in GCPs than in NEPs of Non-IR and IR mice, and there were no significant changes after irradiation (Fig. 1E,F). Consistent with the antibody staining and qRT-PCR results, analysis of RNA-seq data from Nestin-CFP+ NEPs isolated by FACS from P5 IR and Non-IR mice (Wojcinski et al., 2017) showed the *Yap* and *Taz* transcripts were present in NEPs, with no significant changes after irradiation and lower numbers of reads for *Taz* (Table. S1). The DNA-binding TEAD family transcription factors *Tead1* and *Tead2* transcripts were abundant while *Tead3* was minimal (Table. S1), indicating that TEAD1/2 are the main binding partners for YAP/TAZ in neonatal NEPs. The Hippo target gene *Birc5* was present and appears to be slightly upregulated in irradiated NEPs, but a second target gene *Ctgf* had little expression (Table. S1). Together, these data demonstrate that the cell type that predominantly expresses *Yap* and *Taz* is NEPs, and thus YAP and TAZ could play a role in the regeneration of the EGL by NEPs following irradiation.

### Loss of YAP in the NEP lineage hampers postnatal cerebellum regeneration

Given the potential role of YAP in NEPs for CB regeneration, we determined whether *Yap* is required for injury-induced regeneration of the CB by mutating *Yap* specifically in NEPs at P0 using a mosaic mutant analysis approach (MASTR, mosaic mutant analysis with spatial and temporal control of recombination) (Lao et al., 2012; Wojcinski et al., 2017) (Fig. 2A). An inducible FLP site-specific recombinase expressed from a Nestin transgene was used to induce sustained expression of a protein fusion between GFP and CRE (referred to as GFPcre) following injection of tamoxifen (Tm), which then induces recombination of a *Yap* floxed allele (Reginensi et al., 2013), resulting in visualization of the mutant cells and their descendants based on GFP expression (Fig. 2A). Tm was administered to *Nes-FlpoER/+;R26^FSF-GFPcre/+^;Yap^flox/flox^* mice (*Nes-mYap* cKOs) and controls (*Nes-FlpoER/+;Yap^flox/flox^*, or *R26^FSF−^GFPcre/+;Yap^flox/flox^*, or *Yap^flox/flox^*) at P0, X-ray irradiation was conducted at P1, and the size of CB was measured at P30 (Fig. 2B). Notably, *Nes-mYap* cKOs showed a reduction in CB size after irradiation compared to IR controls (Fig. 2E,F; Fig 2S1A). The size of the CB (area of the midline) of IR mutants was reduced to 41.3 ± 2.10% of Non-IR mutants compared to 50.4 ± 2.75% for IR controls compared to Non-IR controls (Fig. 2G). Even more prominent was a significant reduction in the area of the IGL and ratio of IGL/CB area in *Nes-mYap* cKOs compared to controls following irradiation (Fig. 2H-M; Fig. 2S1B,C). Furthermore, as is seen in mutants with a depleted IGL the Calbindin+ Purkinje cells failed to form a single cell layer in the *Nes-mYap* cKOs; instead, the individual Purkinje cells were more disorganized and dispersed throughout the entire WM-IGL-ML layers than in controls (Fig. 2S1D-G,D’-G’). The GFAP+ Bergmann glial fibers also appeared more disorganized in the *Nes-mYap* cKO cerebella (Fig. 2S1H-K, H’-K’). Surprisingly, when the areas of the CB and IGL were measured at P12 and P16, we found that IR *Nes-mYap* cKOs and controls had similar IGL/CB ratios at P12 but a significant reduction at P16 (Fig. 2S2). Together, these data indicate a crucial role of YAP in NEPs at a late stage in the recovery of postnatal CB growth after irradiation-induced injury to the EGL.

**Figure 2.**
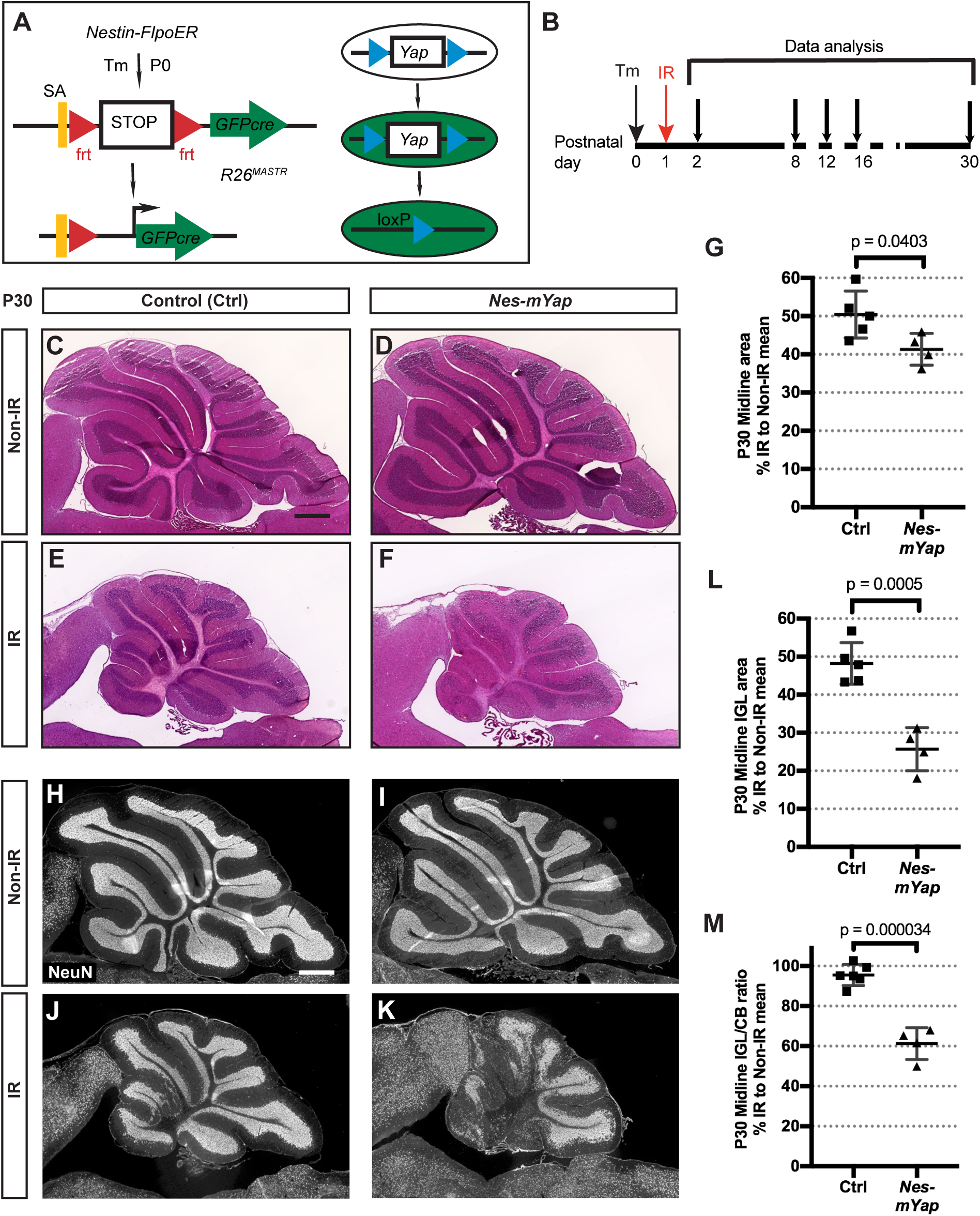
Mutation of *Yap* in the NEP lineage hampers postnatal cerebellum regeneration. (A) Schematic of the MASTR mosaic mutant technique. (B) Schematic showing experimental design. (C-F) H&E staining of midsagittal sections of the CB from Non-IR and IR *Nes-mYap* cKO (Non-IR, n = 6; IR, n = 4) and control mice (Non-IR, n = 4; IR, n = 5) at P30. Scale bar, 500 µm. (G) Graph of the areas of midline sections of the cerebella of IR mice as a percentage of Non-IR animals of the same genotype. (H-K) IF detection of NeuN on midsagittal sections of the CB from Non-IR and IR *Nes-mYap* cKOs and controls at P30. Scale bar, 500 µm. (L-M) Graphs of the IGL areas (L) and IGL/CB ratios (M) of midline cerebellar sections of IR *Nes-mYap* cKOs as a percentage of Non-IR mice of the same genotype (Non-IR, n = 6; IR, n = 4) and controls (Non-IR, n = 4; IR, n = 5) at P30. Data are presented as mean ± S.D., and statistical analysis by unpaired t test. Each data point represents one animal, and is calculated as the value for each IR mouse (average of 3 sections) divided by the mean for all the Non-IR mice of the same genotype, X 100.

### Loss of YAP in NEPs results in an increase in cell death one day following irradiation at P1

Since over-expression of YAP in GCPs was shown to protect that cells from irradiation-induced cell death in culture (Fernandez et al., 2012), we examined whether YAP plays a role in maintaining the survival of NEPs in the PCL following irradiation. As expected, cell death (TUNEL+ particles) was minimal in the CB of Non-IR *Nes-mYap* cKOs and controls given Tm at P0, and cell death was prominent in the EGL of IR mice (Fig. 3A-E). Interestingly, *Nes-mYap* cKOs showed a 1.6-fold increase in the density of TUNEL+ particles within the PCL compared to IR controls (Fig. 3D,E,F). This result indicates that YAP protects NEPs from apoptosis following irradiation.

**Figure 3.**
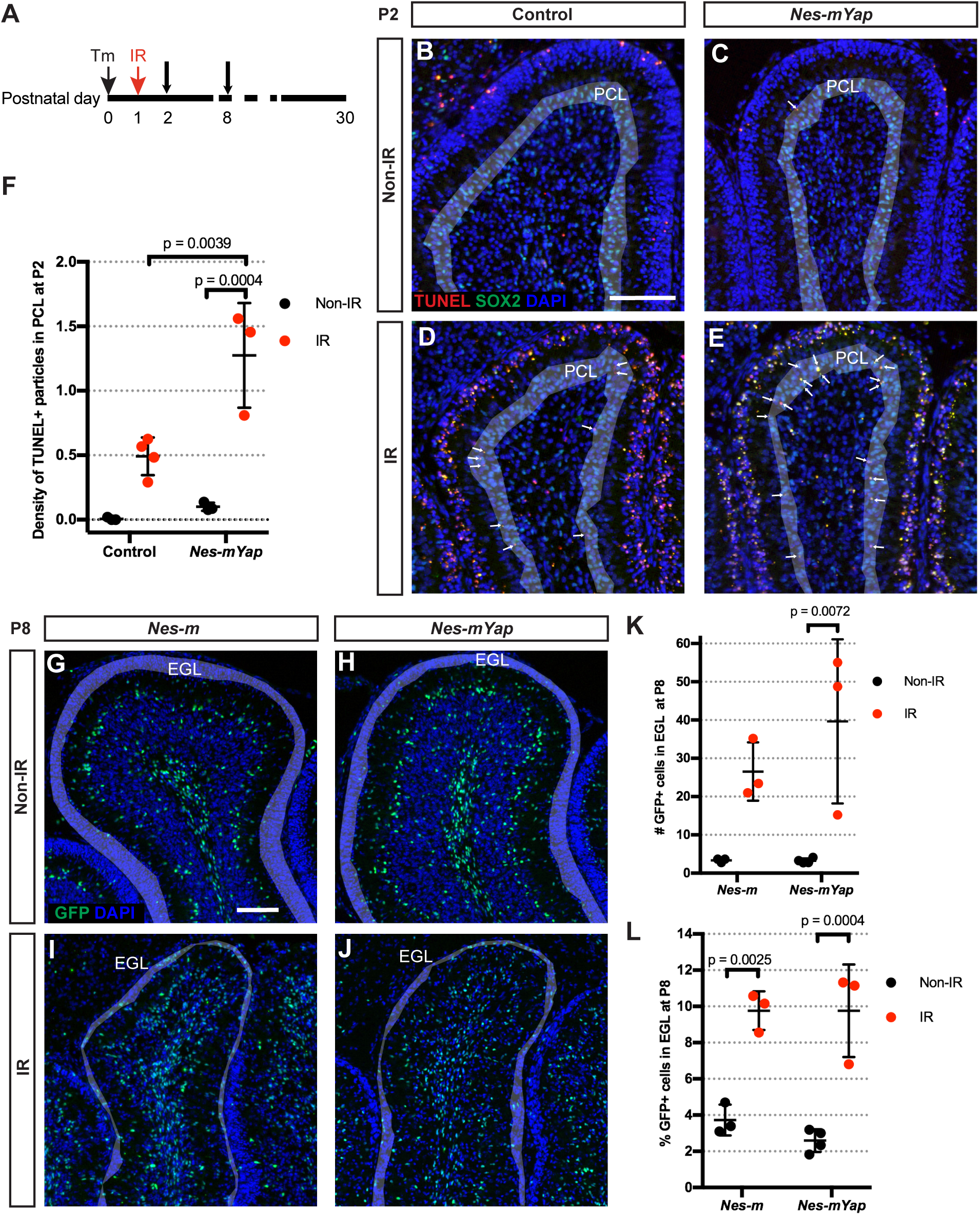
Deletion of of *Yap* in NEPs at P0 results in an increase in cell death in the PCL after irradiation-induced injury at P1. (A) Schematic showing experimental design. (B-E) Representative images from lobule 4/5 showing IF staining for TUNEL, SOX2, and DAPI on midsagittal sections from IR and Non-IR *Nes-mYap* cKOs and controls one day after IR at P1. Grey shadow highlights the PCL of the lobule. Arrows indicate TUNEL+ particles in the PCL. Scale bar, 100 µm. (F) Graph of the densities of TUNEL+ particles in the PCL of the midline CB from IR and Non-IR *Nes-mYap* cKOs (Non-IR, n = 3; IR, n = 3) and controls (Non-IR, n = 3; IR, n = 3) at P2. (G-J) Representative images from lobule 4/5 showing IF staining of GFP and DAPI on midsagittal CB sections from IR and Non-IR *Nes-m* controls and *Nes-mYap* cKOs at P8. Grey shadow highlights the EGL of the lobule. Scale bar, 100 µm. (K-L) Graphs of the numbers of GFP+ cells normalized to total area measured in the EGL (K) and the percentages of GFP+ cells in the EGL among the total number of GFP+ cells (L) from *Nes-m* controls (Non-IR, n = 3; IR, n = 3) and *Nes-mYap* cKOs (Non-IR, n = 4; IR, n = 3) at P8. Data are presented as mean ± S.D., and statistical analysis by two-way ANOVA. Each data point represents one animal.

### YAP is dispensable for the migration of NEPs into the EGL following irradiation-induced injury at P1

Given that the area of the cerebella of *Nes-mYap* cKOs at P12 was similar to Non-IR controls, we examined whether YAP is required for migration of NEPs into the EGL. The distribution of GFP+ cells derived from NEPs labeled at P0 (and also mutated for *Yap* in *Nes-mYap* cKOs) in the different layers of Lobule 4/5 was quantified in both *Nes-mYap* cKOs and controls (*Nes-FlpoER/+;R26^FSF-GFPcre/+^* or *Nes-m*) at P8 (Fig. 3A). Consistent with the initial recovery of CB area in *Nes-mYap* cKOs compared to IR controls, both the number and the percentage of GFP+ cells in the EGL of IR mice was similar between the two genotypes and much greater than in Non-IR mice (Fig. 3G-L). In addition, the numbers and percentages of NEPs in the molecular layer (ML) and IGL+WM were similar between genotypes in the IR condition, but as expected the number of NEPs in the PCL was increased after irradiation (Fig. 3S). These results indicate that the initial responses of NEPs to irradiation do not depend on YAP, including the expansion of the PCL-NEP population and migration of NEPs from the PCL into the EGL to repopulate the GCPs.

### YAP regulates differentiation of NEPs during normal CB development, and the requirement is over-ridden following irradiation

We next examined whether *Yap* is required for the differentiation of NEPs during development of the CB, by determining the distribution of the descendants of NEPs labeled at P0 into the cerebellar layers at P8 in *Nes-mYap* cKOs and *Nes-m* controls. Quantification of the GFPcre+ NEP-derived cells in lobule 4/5 of P8 demonstrated a significant increase in the number of GFP+ cells in the ML normalized to the area of the lobule analyzed, as well as the percentage of cells in that layer in *Nes-mYap* cKOs compared to *Nes-m* controls (Fig. 4A-E; Fig. 4S1A,B), and possibly an increase in the number of mutant cells present at P8 (Fig. 4C; Fig. 4S1A,B). There was a concomitant decrease in the percentage of cells in the IGL+WM layers, but not the total number of cells, indicating that the significant change in *Nes-mYap* cKOs is an increase in the production of NEP-derived cells that populate the ML (Fig. 4E; Fig. 4S1A,B). Consistent with the GFP+ cells in the ML being the expected interneurons produced by WM-NEPs, there was also a significant increase in the number of cells in the ML that expressed PAX2, a marker of differentiating interneurons (Fig. 2F). The number of PAX2+ cells in the IGL+WM also might be increased, consistent with an overall increase in production of interneurons in the absence of *Yap* (Fig. 4G; Fig. 4S1C,D). Quantification of S100β+ astrocytes among the GFP+ populations in lobule 4/5 revealed an increase in the number of astrocytes in the PCL (Fig. 4H; Fig. 4S1E,F). These data reveal that YAP normally plays a role in attenuating production of interneurons and astrocytes.

**Figure 4.**
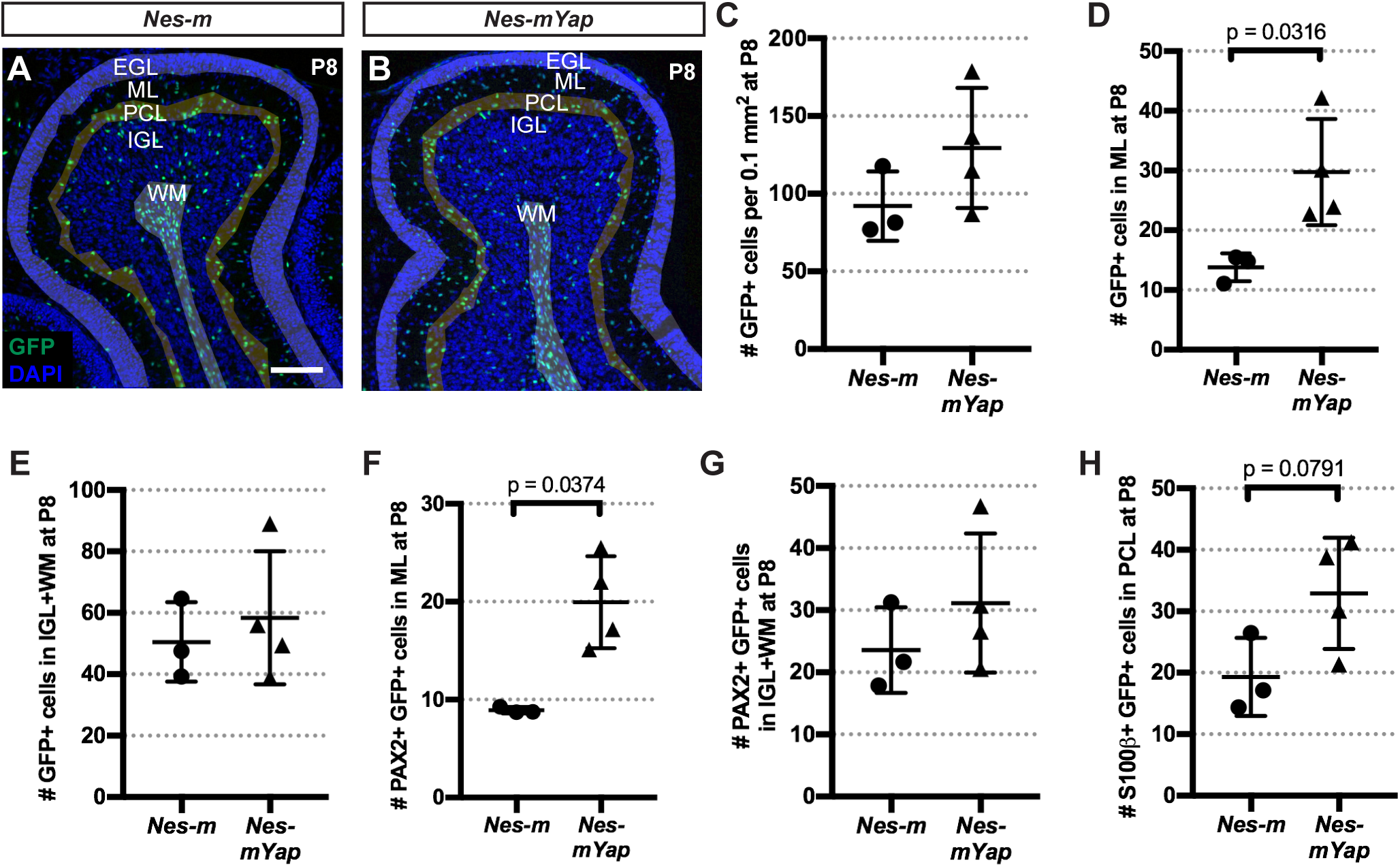
YAP promotes the differentiation of NEPs during normal postnatal CB development. (A-B) Representative images from lobule 4/5 showing IF staining of GFP and DAPI on midsagittal sections from a *Nes-m* control and a *Nes-mYap* cKO at P8. Shading indicates the partitioning of the different layers (EGL, ML, PCL, IGL and WM) within the lobule. Scale bar, 100 µm. (C-H) Graphs of the numbers of GFP+ cells in all the layers (C) or in the ML (D) or IGL+WM (E), and the numbers of PAX2+ GFP+ cells in the ML (F) and IGL+WM (G), and the numbers of S100β+ GFP+ cells in the PCL (H), per 0.1 mm^2^ of the total area analyzed in lobule 4/5 from *Nes-m* controls (n = 3) and *Nes-mYap* cKOs (n = 4) at P8. (C) P = 0.1996; (E) P = 0.6065; (G) P = 0.3548. Statistical analysis is conducted by unpaired t test. Data are presented as mean ± standard deviation (S.D.), and each data point represents one animal.

Given that we did not observe a change in the distribution of *Yap* mutant GFP+ cells in the layers of the P8 CB after irradiation, we next examined whether the loss of YAP alters the differentiation of NEPs into interneurons (PAX2+) and astrocytes (S100β+) during irradiation-induced recovery of the CB, since both were increased in Non-IR mutants. Although the normalized number of PAX2+ interneurons in lobule 4/5 at P8 showed a significant difference between *Yap* mutant and control IR mice, no difference was observed in the number or percentage of PAX2+ cells in a specific layer (Fig. 4S2A,B). Additionally, there was no difference in the production of S100β+ astrocytes between *Yap* mutant and control mice after irradiation (Fig. 4S2C,D). These results indicate that the requirement for YAP in differentiation of NEPs is over-ridden when the EGL of the CB is injured.

### Loss of YAP results in an increase in cell death in the EGL at P12 following irradiation-induced injury at P1

We next analyzed the CB at P12, when the EGL is normally diminishing due to increased production of granule neurons. Given the reduction of the IGL in mutants compared to control IR mice at P30, we asked whether YAP plays a role in cell survival of GCPs. Strikingly, TUNEL staining in the EGL revealed a significant increase in the density of TUNEL+ particles within the EGL (number of TUNEL+ particles per 0.1mm^2^) of *Nes-mYap* cKOs compared to controls after irradiation (Fig. 5). In contrast, the density of TUNEL+ particles within the EGL at P8 was similar between controls and mutants, consistent with the similar distribution of GFP+ cells between the cell layers (Fig. 5S). These results indicate that YAP is required for GCPs to maintain cell survival in the EGL during injury-induced recovery.

**Figure 5.**
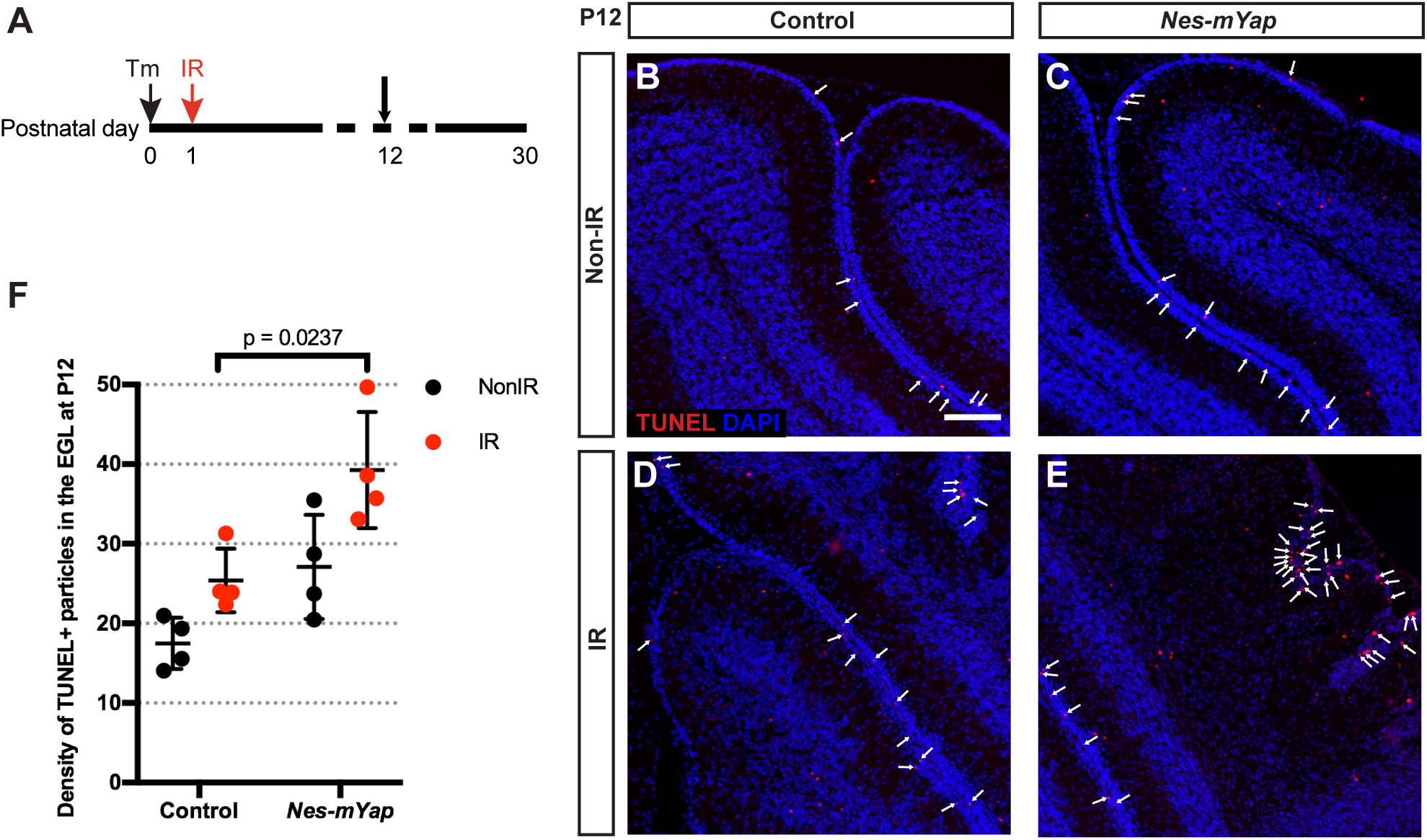
Mutation of *Yap* in NEPs at P0 results in more cell death in the EGL at P12 after irradiation-induced EGL injury at P1. (A) Schematic showing experimental design. (B-E) Representative images showing IF staining for TUNEL and DAPI on midsagittal sections from IR and Non-IR *Nes-mYap* cKOs and controls at P12. Arrows indicate TUNEL+ particles in the EGL. Scale bar, 100 µm. (F) Graph of the densities of TUNEL+ particles in the EGL (the number of TUNEL+ particles per 0.1 mm^2^ of EGL area) in midline CB sections from *Nes-mYap* cKOs (Non-IR, n = 3; IR, n = 4) and controls (Non-IR, n = 4; IR, n = 4) at P12. Data are presented as mean ± S.D., and statistical analysis by two-way ANOVA. Each data point represents one animal.

### YAP is not required in GCPs for cerebellar growth during normal development

Given that when *Yap* is deleted in NEPs there is an increase in cells death of GCPs at P12, one possible explanation is that YAP is required for GCP proliferation/differentiation or survival. In order to determine whether YAP plays a role in GCPs during normal development of the CB, we generated two mutants. First, we deleted *Yap* from the ATOH1-expressing rhombic lip lineage when GCPs are generated in the embryo and analyzed the size of the CB at P30. Consistent with the weak expression of *Yap* in GCPs (Fig. 1A-C), P30 *Atoh1-Cre/+;Yap^flox/flox^ (Atoh1-Yap* cKO) mice showed no reduction in the area of the midline CB (Fig. 6A,B), or the IGL area or IGL/CB ratio compared to *Yap^flox/flox^* littermate controls (Fig. 6C-G). This lack of a growth phenotype is similar to mice lacking *Yap* in NEPs (Fig. 2C,D), showing that loss of YAP in the *Atoh1*-lineage does not have a major effect on growth of the CB during development.

**Figure 6.**
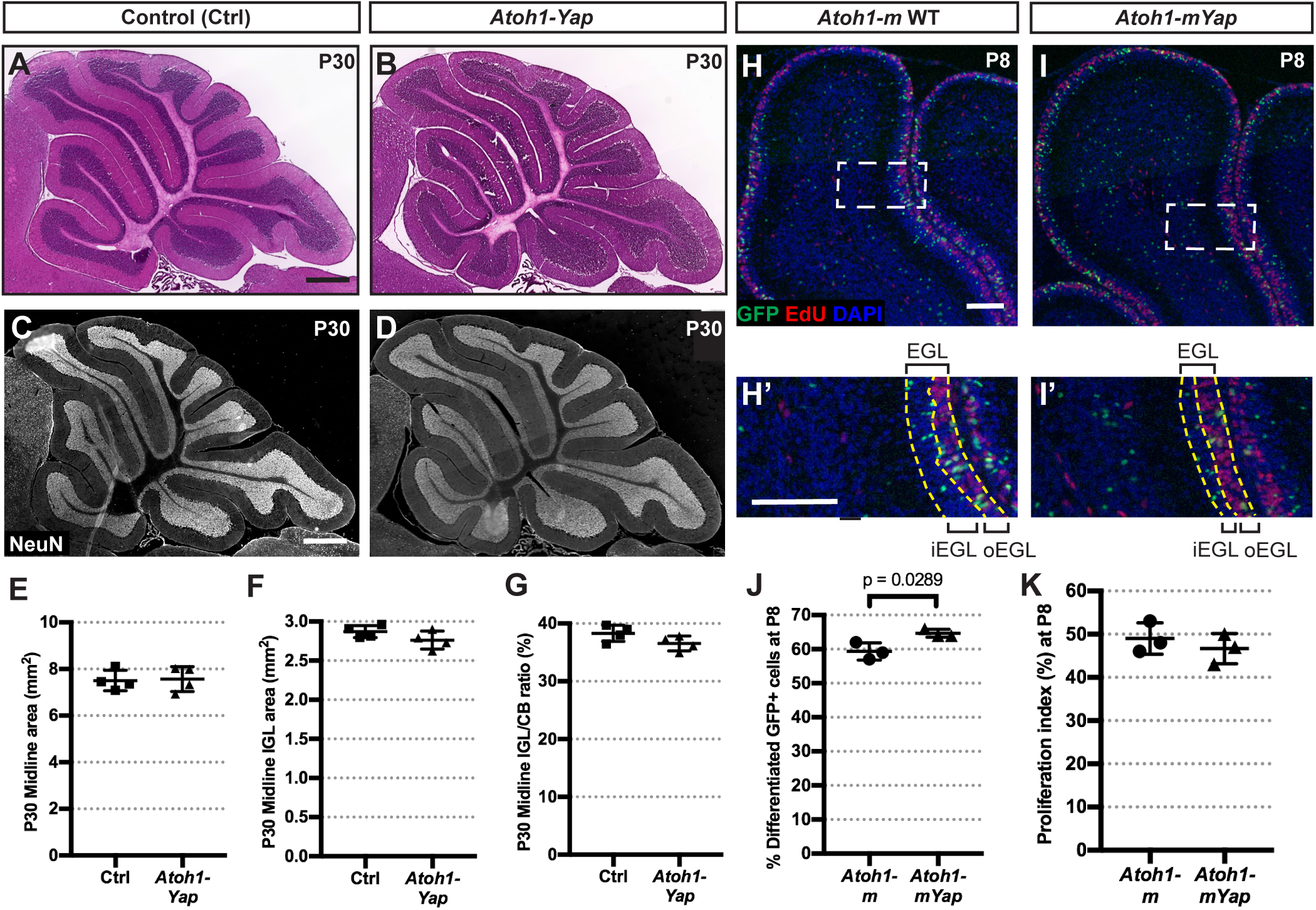
YAP promotes differentiation of GCPs during postnatal development of the CB. (A-D) H&E staining (A-B) and IF detection of the granule cell marker NeuN (C-D) on midsagittal sections of cerebella from IR and Non-IR *Atoh1-Yap* cKOs and control (Ctrl) mice at P30. Scale bars, 500 µm. (E-G) Graphs of the areas of the midline of the CB (E), the areas of the IGL (F), and IGL/CB area ratios (G) in IR *Atoh1-Yap* cKO (n = 4) and littermate controls (n = 4) at P30 as a ratio of Non-IR mice of the same genotype. (E) p = 0.8554; (F) p = 0.1727; (G) p = 0.1155. (H-I) Representative images from lobule 4/5 showing IF staining for GFP, EdU, and DAPI on midsagittal sections from an *Atoh1-m* control and *Atoh1-mYap* cKO at P8. (H’-I’) Magnification of areas within dotted lines in H-I. Yellow dashed lines indicate the EGL and the border of the outer- and inner-EGL. Scale bars, 100 µm. (J-K) Graphs of the percentages of GFP+ differentiating cells (in all layers except the oEGL) among all GFP+ cells (J) and proliferation indices (G) in the midline of the CB in *Atoh1-m* controls (n = 3) and *Atoh1-mYap* cKOs (n = 3) at P8. (K) P = 0.4670. Data are presented as mean ± S.D., and statistical analysis by unpaired t test. Each data point represents one animal.

We next used the MASTR mosaic mutant approach with an *Atoh1-FlpoER* transgene (Wojcinski et al., 2019) as a sensitive assay for a mild change in differentiation of GCPs when *Yap* is removed. Tm was administered at P0 to *Atoh1-FlpoER/+;R26^FSF-GFPcre/+^;Yap^flox/flox^ (Atoh1-mYap* cKO) mice and controls (*Atoh1-FlpoER/+; R26^FSF-GFPcre/+^* or *Atoh1-m*), and then EdU was injected 1 hour prior to sacrifice at P8. The EdU+ outer EGL (oEGL) consists of actively proliferating GCPs, and the percentage of post-mitotic GFP+ GCs in the EdU-negative “inner layers” including the inner EGL (iEGL), ML, IGL, and WM was determined amongst all GFP+ cells (Fig. 6H-I, H’-I’). Interestingly, we found that the percentage of GFP+ cells in the “inner layers” was slightly but significantly higher in *Atoh1-mYap* cKOs compared to *Atoh1-m* controls (Fig. 6J), indicating an increase in differentiation, or decrease in self-renewal of GCPs. The proliferation index (percent of EdU+ GFP+ cells among all GFP+ cells in the oEGL) was similar between GFP+ *Atoh1-mYap* cKO cells and *Atoh1-m* control cells (Fig. 6K), showing that *Yap* does not regulate the proliferation rate. Taken together, these data indicate YAP plays a minor but negative role on differentiation of GCPs into granule neurons. Given this role of *Yap*, all be it mild, in promoting GCP self-renewal one might have expected a growth defect in *Atoh1-Yap* cKOs. One possibility is that this phenotype is only expressed in a mosaic situation, where there is competition between the scattered mutant cells and their surrounding wild type neighbors.

### Loss of *Taz* in *Yap* mutant NEPs does not abrogate recovery after irradiation

Since TAZ and YAP have been found to have both similar and distinct requirements in the development or regeneration of various organs (Deng et al., 2016; Hossain et al., 2007; Makita et al., 2008; Morin-Kensicki et al., 2006; Reginensi et al., 2013; Tian et al., 2007), and we detected nuclear TAZ in NEPs (Fig. 1CD,C’-D’, C”-D”), we asked whether TAZ contributes to the regenerative response following irradiation in *Nes-mYap* cKOs. We first tested any requirement for *Taz* in development of the CB and replenishment of the CB after irradiation by administering Tm at P0 to *Nes-FlpoER/+;R26^FSF-GFPcre/+^;Taz^flox/flox^* mice (*Nes-mTaz* cKOs) and controls (*Nes-FlpoER/+;Taz^flox/flox^*, or *R26^FSF-GFPcre/+;Tazflox/flox^*, or *Taz^flox/flox^*). Similar to *Nes-mYap* cKOs, conditional deletion of *Taz* did not alter cerebellar size at P30 (Fig. 7; Fig. 7S). However, unlike *Nes-mYap* cKOs, *Nes-mTaz* cKOs recovered almost as well as control mice after irradiation (Fig. 7; Fig. 7S). We next ablated both *Yap* and *Taz* from NEPs using *Nes-FlpoER/+;R26^FSF-GFPcre/+^;Yap^flox/flox^;Taz^flox/flox^* mice (*Nes-mYapTaz* cKOs) and controls (*Nes-FlpoER/+;Yapflox/flox;Tazflox/flox*, or *R26FSF-GFPcre/+;Yapflox/flox;Tazflox/flox*, or *Yap^flox/flox^;Taz^flox/flox^*) given Tm at P0. Surprisingly, ablation of both *Yap* and *Taz* did not result in a worse recovery of the CB after injury compared to *Nes-mYap* cKOs. On the contrary, *Nes-mYapTaz* cKOs had no significant reduction in the area of the midline CB or IGL, although they had a small but significant reduction in the IGL/CB ratio compared to controls (Fig. 8; Fig. 8S1A-C). The apparent better recovery of *Nes-mYapTaz* cKOs compared to *Nes-mYap* cKOs could indicate that loss of *Taz* partially rescues the poor late recovery observed in *Nes-mYap* cKOs (Fig. 8S1D-F).

**Figure 7.**
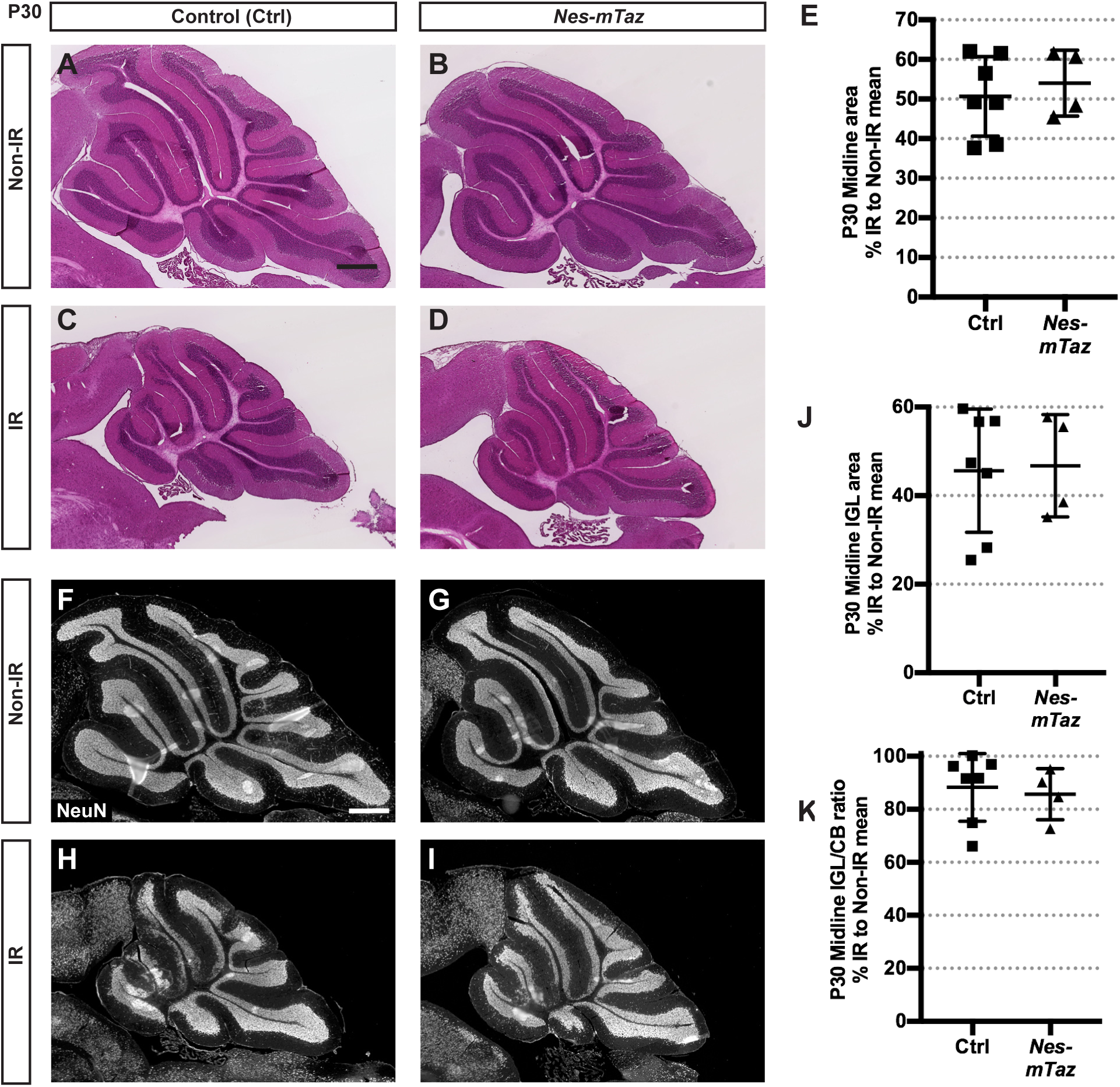
Mutation of *Taz* in NEPs at P0 does not affect cerebellar growth during development and regeneration after irradiation at P1. (A-D) H&E staining of midsagittal sections of cerebella from Non-IR and IR of *Nes-mTaz* cKOs and controls at P30. Scale bars, 500 µm. (E) Graph of the areas of the CB of IR animals as a percentage of Non-IR mice of the same genotype in midline sections. p = 0.5866. (F-I) IF detection of NeuN on midsagittal sections of the CB from Non-IR and IR *Nes-mTaz* cKOs and controls at P30. Scale bar, 500 µm. (J,K) Graphs of the IGL areas (J) and IGL/CB ratios (K) of IR *Nes-mTaz* cKOs (Non-IR, n = 6; IR, n = 4) and controls (Non-IR, n = 4; IR, n = 7) at P30 as a percentage of Non-IR same genotype. (J) p = 0.8945; (K) p = 0.7335. Data are presented as mean ± S.D., and statistical analysis by unpaired t test. Each data point represents one animal, and is calculated using each IR measurement divided by the mean of Non-IR mice of the same genotype, X 100.

**Figure 8.**
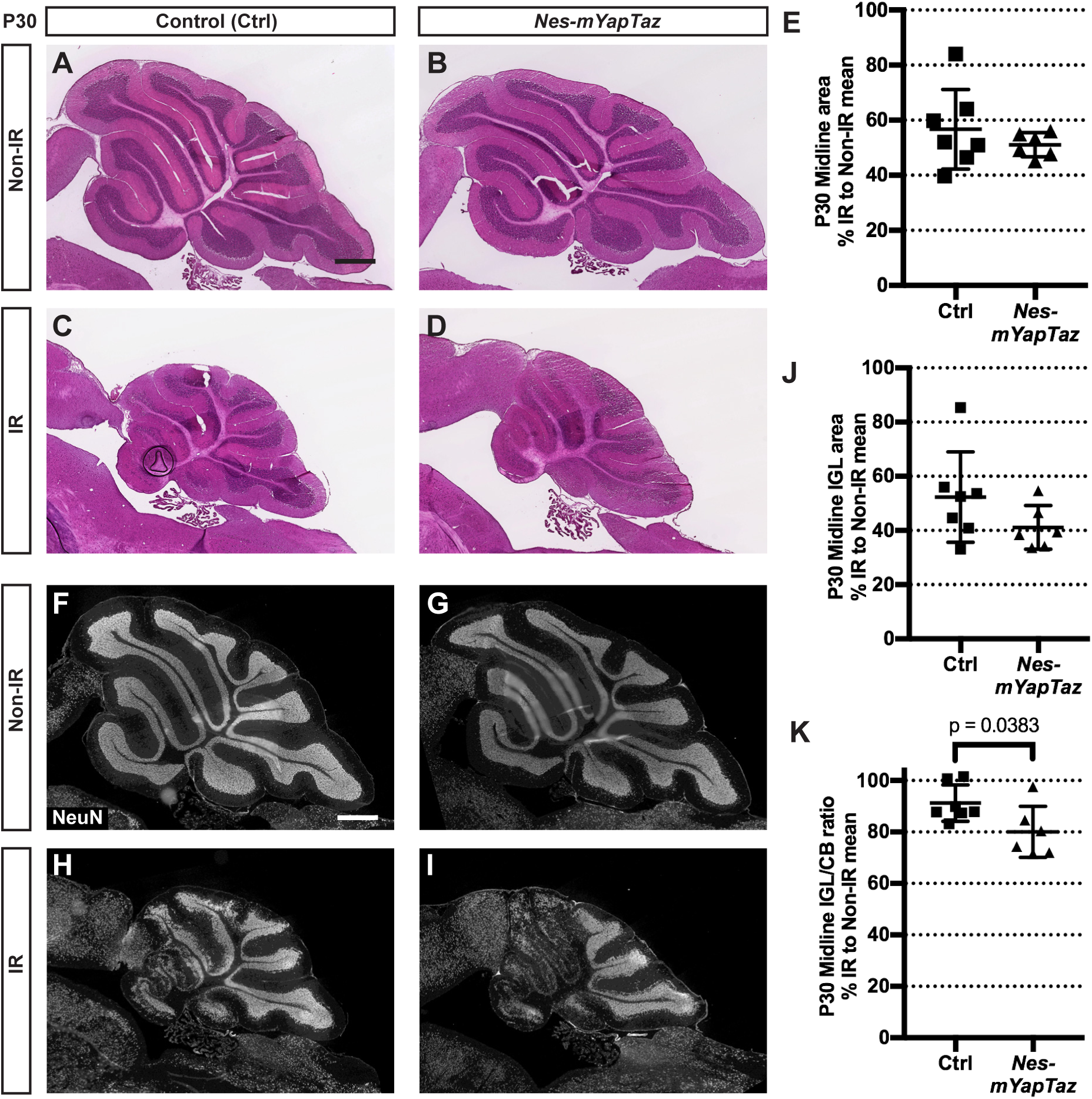
Loss of *Taz* in NEPs lacking *Yap* at P0 does not abrogate recovery of the IGL after irradiation at P1. (A-D) H&E staining of midsagittal sections of the CB from Non-IR and IR *Nes-mYapTaz* cKOs and controls at P30. Scale bars, 500 µm. (E) Graph of the midline CB areas of IR mice as a percentage of Non-IR animals of the same genotype. P = 0.3815. (F-I) IF detection of NeuN on midsagittal sections of the CB from Non-IR and IR animals at P30. Scale bar, 500 µm. (J,K) Graphs of the areas of the IGL (J) and IGL/CB ratios (K) of IR mice as a percentage of Non-IR animals of the same genotype on midsaggital sections at P30. *Nes-mYapTaz* cKOs (Non-IR, n = 7; IR, n = 6) and controls (Non-IR, n = 4; IR, n = 7). (J) p = 0.1637; (K) p = 0.0383. Data are presented as mean ± S.D., and statistical analysis by unpaired t test. Each data point represents one animal, and is calculated using the average for each IR mouse divided by the mean of the Non-IR mice of the same genotype, X 100.

In order to examine whether the apparent better recovery in *Nes-mYapTaz* cKOs compared to *Nes-mYap* cKOs correlates with better survival of GCPs in the EGL at P12, we quantified cell death by TUNEL staining of cerebellar sections. Unlike the obvious increase in the density of TUNEL+ particles within the EGL of *Nes-mYap* cKOs compared to controls after irradiation (Fig. 5F), IR *Nes-mYapTaz* cKOs did not have a significant increase in cell death in the EGL compared to controls (Fig. 8S2). Such an attenuated cell death could contribute to the slightly better regeneration observed in *Nes-mYapTaz* cKOs compared to *Nes-mYap* cKOs.

## DISCUSSION

In this study, we demonstrated that YAP has a major requirement for postnatal regeneration of the CB following irradiation. In particular, mutation of *Yap* impairs the recovery of normal cerebellar size and formation of the IGL following irradiation. This defect is accompanied by an increase in cell death in the EGL at P12, following normal initial expansion and migration of NEPs to the EGL in response to the injury. The organizations of Purkinje cells and Bergmann glial fibers were also disrupted in *Yap* mutants. It is possible that this aspect of the phenotype is a secondary result of the failure of the GCPs to generate an IGL in IR *Yap* mutants. Alternatively, *Yap* depletion could result in a cell autonomous defect in Bergamnn glia that is only critical following irradiation, which in turn reduces survival of GCPs and/or hinders the migration of GCPs into the IGL resulting in a depletion of the EGL.

The pivotal role of Hippo signaling in brain development has been extensively studied in *Drosophila*, a model organism in which the Hippo pathway itself was first discovered. Yorkie-null mutant Drosophila display semi-lethality and impaired proliferation and growth of neural stem cells at the larval stage (Ding et al., 2016). In a complementary manner, inhibition of Hippo signaling, through either over-expression of the *Drosophila* YAP homolog Yorkie or loss-of-function mutations in several core kinases that activate Yorkie function, results in increased neuroblast proliferation and substantial brain overgrowth (Poon et al., 2016). In the mouse, genetic ablation of *Yap* in radial glial progenitors leads to only subtle proliferation defects during cortical development (Park et al., 2016), whereas loss of both *Yap* and *Taz* results in premature differentiation of radial glial progenitors coupled with precocious production of neurons (Kong, 2018) or a loss of ependymal cells that line the ventricle in postnatal brain (Liu et al., 2018). Consistent with the subtle requirement for YAP in cortical development (Park et al., 2016), we found that *Yap* ablation from NEPs or GCPs does not significantly reduce the overall size of the adult CB. We nevertheless observed mild increases in differentiation of *Yap* mutant NEPs and GCPs. In particular, removal of *Yap* from NEPs or GCPs in a mosaic manner in the CB between P1 and P8 results in an increase in differentiation of NEPs (PCL-NEPs to produce astrocytes and WM-NEPs to produce interneurons) and GCPs (to produce granule cells) possibly at the expense of self-renewal. YAP seems to have a similar function in progenitor proliferation/differentiation in other organs. For example, YAP over-activation in mouse intestine or chick neural tube leads to expansion of progenitor cells and loss of differentiated cells (Camargo et al., 2007; Cao et al., 2008). It also has been shown that over-expression of YAP in cultured myoblasts inhibits differentiation and formation of myotubes (Watt et al., 2010), and reduced Hippo signaling in mouse epithelial cells results in hyper-proliferation of progenitors and failure of differentiation into multiple epithelial cell types (Lee et al., 2008). Thus, a consistent finding from our current study in the CB and work of others in distinct organs is that YAP modulates a timely transition of neural progenitors from cell proliferation into differentiation.

Several lines of evidence suggest an intersection or coupling of the Hedgehog (HH) and Hippo pathways in *Drosophila* and mice (Akladios et al., 2017; Fernandez et al., 2009; Hsu et al., 2017; Huang and Kalderon, 2014; Lin et al., 2012). For example, in flies elevated Hh signaling (*ptc* mutants) induces Yorkie target gene transcription, whereas additional deletion of Yorkie abolishes the effects of excess Hh signaling on cell proliferation and survival (Huang and Kalderon, 2014). In cultured mouse GCPs, SHH treatment increases the transcription and nuclear localization of YAP protein (Fernandez et al., 2009). Our previous study showed that SHH signaling is required for NEPs to regenerate the mouse CB, particularly at the initial stage of recovery (Wojcinski et al., 2017). Thus it is possible that SHH signaling after irradiation activates YAP function and this promotes survival of NEPs immediately after irradiation, since we observed an increase in cell death after irradiation. However, comparison of *Nes-mSmo* cKO (Wojcinski et al., 2017) and *Nes-mYap* cKO phenotypes after irradiation of the CB reveals additional requirements for SHH signaling that are independent of *Yap*. There is an immediate defect in expansion of NEPs and their migration to the EGL in *Nes-mSmo* cKOs after irradiation (Wojcinski et al., 2017), whereas these cellular responses are intact in *Nes-mYap* cKOs. Taken together, the results provide strong evidence that SHH has critical target genes other than *Yap* that are required for proliferation and migration of NEPs.

YAP and TAZ have often been found to have overlapping functions, consistent with their significant homology and common binding partners, but some recent studies point to distinct roles in addition to overlapping functions of YAP and TAZ that appear to be context dependent. For example, *Yap* null mutant mice are embryonical lethals (Morin-Kensicki et al., 2006), whereas *Taz* nulls only show partial embryonic lethality and mice that survive have lung defects and kidney disease (Hossain et al., 2007; Makita et al., 2008; Tian et al., 2007). It has also been shown that TAZ competes with YAP in an interaction with a regulator protein that modulates the maturation of chondrocytes, a critical step for skeletal development and bone repair (Deng et al., 2016). Similarly we found that *Yap* ablation from NEPs leads to poor recovery of the postnatal CB following acute injury, but deletion of *Taz* does not abrogate regeneration. Double conditional *Yap* and *Taz* mutants instead appear to have a milder phenotype than *Yap* cKOs, indicating that loss of *Taz* partially rescues the *Yap* mutant phenotype. This finding raises the possibility of antagonistic functions between YAP and TAZ in NEPs, as in the bone, compared to overlapping functions under other circumstances. Alternatively, the difference in genetic backgrounds of the single and double mutants could contribute to the degree of recovery of *Yap* mutant NEPs after irradiation. Thus it will be important to dissect the distinct downstream molecular pathways of the two co-activators to reveal their context-dependent functions in the Hippo pathway. Leveraging this knowledge will pave the road for potential therapeutic intervention for cerebellar hypoplasia caused by injury

### Materials and Methods

#### Mice

The following mouse lines were used: *Nes-CFP* (Mignone et al., 2004), *Atoh1-GFP* (Chen et al., 2002), *Nestin-FlpoER* (Wojcinski et al., 2017), *Atoh1-FlpoER* (Wojcinski et al., 2019), *Rosa26^MASTR(frt-STOP-frt-GFPcre)^* (Lao et al., 2012), *Yap^flox/flox^* (Reginensi et al., 2013), and *Atoh1-Cre* (Matei et al., 2005). Tamoxifen (Sigma, T5648) was dissolved in corn oil at 20 mg/mL and a single dose of 200 µg/g was administered to P0 animals by subcutaneous injection. Mouse husbandry and all experiments were performed in accordance with MSKCC IACUC-approved protocols.

#### Irradiation

P1 mice were anesthetized by hypothermia and received a single dose of 4 Gy irradiation in an X-RAD 225Cx (Precision X-ray) Microirradiator in the MSKCC Small-Animal Imaging Core Facility. A 5-mm diameter collimator was used to target the CB from the left side of the animal.

#### Tissue Processing

For animals younger than P4, brains were dissected out and fixed in 4% paraformaldehyde overnight at 4°C. Animals P4-30 were anesthetized and transcardially perfused with PBS followed by chilled 4% paraformaldehyde. Brains were harvested and post-fixed overnight and cryoprotected in 30% sucrose before freezing in Cryo-OCT. Frozen brains were sectioned at 12 µm on a cryostat, and sagittal sections of the midline CB were used for all analyses.

#### Immunofluorescent staining

Cryosections were stained overnight at 4 °C with the following primary antibodies: mouse anti-YAP (Abcam, AB56701), rabbit anti-TAZ (Santa Cruz, sc-48805), rat anti-GFP (1:1,000; Nacalai Tesque; 0440484), mouse anti-NeuN (Millipore, MAB377), rabbit anti-Calbindin D-28K (Swant, CB38), rabbit anti-GFAP (Dako, Z0334), rabbit anti-S100β (Dako, Z0311), rabbit anti-PAX2 (Invitrogen, 71600), and goat anti-SOX2 (R&D System, AF2018). Secondary antibodies for double labeling were donkey anti-species conjugated with Alexa Fluor 488 or 555 (1:1,000; Molecular Probes). Nuclei were counterstained with Hoechst 33258 (Invitrogen, H3569).

#### Microscopy

Images were collected either on a DM6000 Leica microscope using Zen software (Zeiss), or a NanoZoomer 2.0 HT slide scanner (Hamamatsu Photonics) using NDP.scan software. All images were taken with 20X objectives, and processed using NDP.view2 and Photoshop softwares.

#### Flow Cytometry

Cerebella of *Atoh1-GFP* and *Nestin-CFP* mice were dissected out under the dissection microscope. Tissues were digested by Trypsin/DNase, and then subject to FACS (fluorescence activated cell sorting) to isolate GFP+ or CFP+ cells. RNA was extracted from GFP+ or CFP+ cells of individual cerebella and then subject to qRT-PCR analysis.

#### qRT-PCR

RNA was isolated from FACS-isolated GFP+ cells from P4 *Atoh1-GFP* mice and FACS-isolated CFP+ cells from *Nestin-CFP* mice using a miRNeasyMicro Kit (Qiagen) according to the manufacturer’s protocol. cDNA was prepared using iScript cDNA synthesis kit (Bio-Rad). qRT-PCR was performed using PowerUp Sybr Green Master Mix (Applied Biosystems). Fold changes in expression were calculated using the ΔΔCt method. The *Gapdh* gene was used to normalize the results. The following primer pairs were used: *Yap* forward 5′-ACCCTCGTTTTGCCATGAAC-3′, *Yap* reverse 5′-TGTGCTGGGATTGATATTCCGTA-3′, *Gapdh* forward 5′-CCAAGGTGTCCGTCGTGGATCT-3′, and *Gapdh* reverse 5′-GTTGAAGTCGCAGGAGACAACC-3’.

#### Quantification and statistical analysis

ImageJ software was used to measure the area (mm^2^) of the CB on sections near the midline. For all IF staining, cell counts were obtained using Stereo Investigator Software. Three sections per animal and at least three animals were analyzed for quantifications. Statistical analyses were performed using Prism software (GraphPad) and significance was determined at P < 0.05. All statistical analyses were two-tailed. For two-group comparisons with equal variance as determined by the F-test, an unpaired Student’s t test was used. For comparisons among four independent groups with equal variance, an unpaired one-way ANOVA was used. For comparisons between groups that are split with two independent variables, an unpaired two-way ANOVA was used. P values and degrees of freedom are given in the figures and legends. Data are presented as mean ± S.D. (standard deviation). No statistical methods were used to predetermine the sample size, but our sample sizes are similar to those generally employed in the field. No randomization was used. Data collection and analysis were not performed blind to the conditions of the experiments.

#### EdU (5-ethynyl-2’-deoxyuridine) Injection and Staining

For assessing cell proliferation, EdU (Invitrogen, E10187) was given at 100 mg/g by i.p. injection 1 h before euthanasia. Click-it EdU assay with Sulfo-Cyanine5 azide (Lumiprobe corporation, A3330) was used according to the protocol of the manufacturer.

#### TUNEL Staining

For TUNEL staining, slides were permeabilized with 0.5% TritonX-100, pre-incubated with Tdt buffer (30 mM Tris•HCl, 140 mM sodium cacodylate and 1 mM CoCl^2^) for 15 min at room temperature, and incubated for I h at 37 °C in TUNEL reaction solution containing Terminal Transferase (Roche, 03333574001) and Digoxigenin-11-dUTP (Sigma-Aldrich, 11093088910). Slides were then incubated with anti-digoxigenin-rhodamine (Sigma-Aldrich, 11207750910) for 1 h.

## Acknowledgements

We thank Dr. Alexandre Wojcinski for insightful suggestions on experimental design and interpretation of results, and Dr. I-Li Tan for performing a RT-Q-PCR experiment. We are grateful to Daniel Stephen and Zhimin Lao for technical assistance. We thank all the Joyner lab members for their helpful comments and discussions. We also thank the Flow Cytometry core and the Center for Comparative Medicine and Pathology of MSKCC for outstanding technical support. We gratefully acknowledge P. Zanzonico for his help with mouse irradiation; Q. Chen and the MSKCC Small-Animal Imaging Core Facility for technical assistance; and a Shared Resources Grant from the MSKCC Geoffrey Beene Cancer Research Center, which provided funding support for the purchase of the XRad 225Cx Microirradiator.

## Competing interests

The authors declare no competing or financial interests.

## Author contributions

A.L.J. helped design the study, interpret the data and write the manuscript, and supervised the project. Z.Y. helped design the study, and analyze and interpret the data. Z.Y. also performed most of the experiments and wrote a first draft of the manuscript.

## Funding

This work was supported by a grant from the NIH (R01 NS092096) to A.L.J and a National Cancer Institute Cancer Center Support Grant (P30 CA008748-48).

