## Supplemental Data for "YAP is involved in replenishment of granule cell progenitors following injury to the neonatal cerebellum"

**Table S1**. Read number of Hippo pathway genes from RNA-sequencing of Non-IR and IR CFP+ NEPs isolated from P5 *Nestin-CFP* mice.

| **Gene** | **Non-IR NEPs** | **IR NEPs** | **Unadjusted p value** |
|---|---|---|---|
| *Yap* | 3203 | 3051 | 0.88 |
| *Taz* | 2129 | 2211 | 0.73 |
| *Tead1* | 8018 | 8137 | 0.91 |
| *Tead2* | 7219 | 7587 | 0.64 |
| *Tead3* | 648 | 490 | 0.08 |
| *Ctgf* | 72 | 86 | 0.50 |
| *Birc5* | 2687 | 3125 | 0.19 |
| *Gapdh* | 2081 | 2212 | 0.60 |

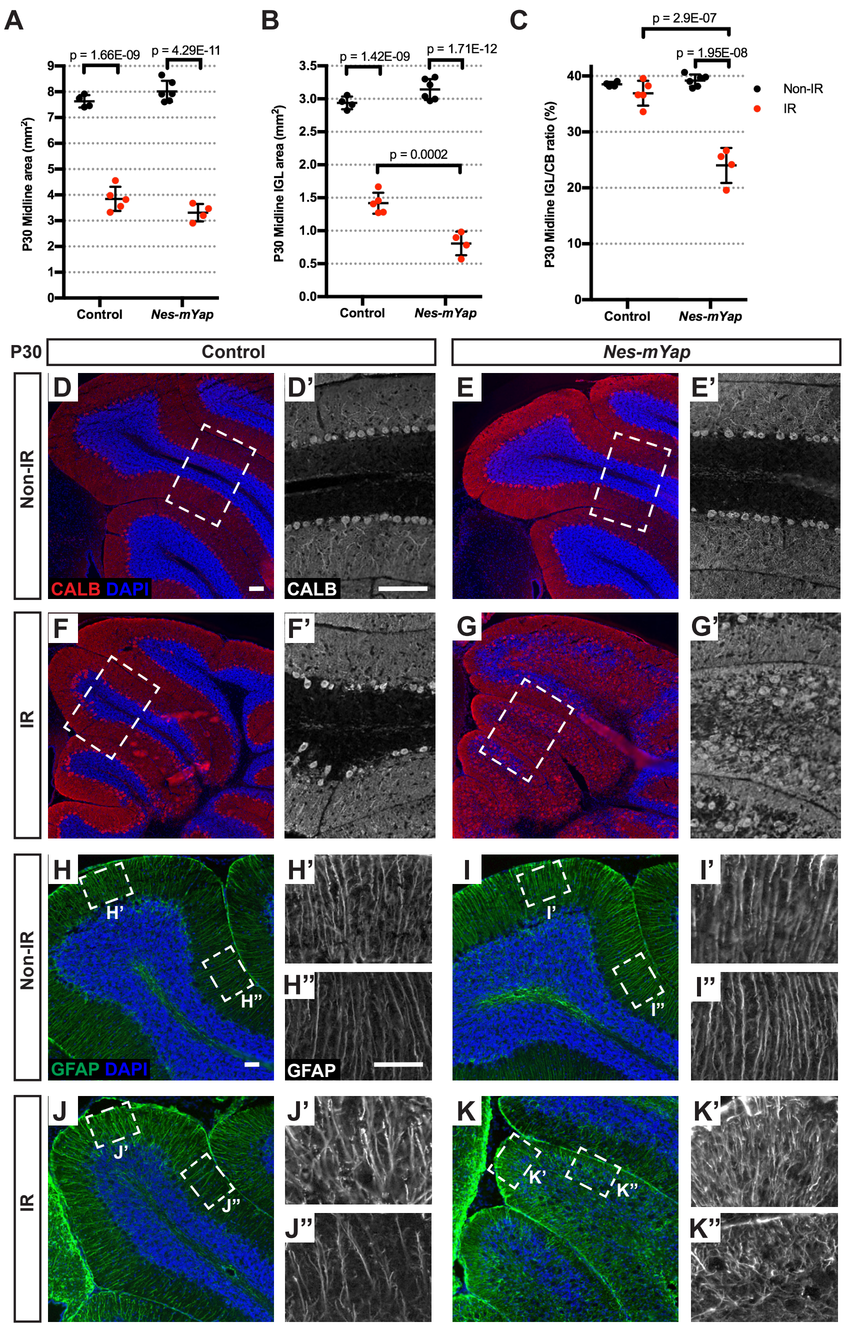


**Figure 2S1. Deletion of of *Yap* in NEPs hinders injury-induced regeneration of the CB and disrupts the layered cytoarchitecture.**
(A-C) Graphs of the midline cerebllar areas (A), IGL areas (B), and IGL/CB ratios (C) from *Nes-mYap* cKOs (Non-IR, n = 6; IR, n = 4) and controls (Non-IR, n = 4; IR, n = 5) at P30. Data are presented as mean ± S.D., and statistical analysis by two-way ANOVA. Each data point represents one animal. (D-K) Representative images from lobule 4/5 showing IF staining of Calbindin (CALB) (D-G) and GFAP (H-K) on midsagittal sections from *Nes-mYap* cKOs and controls at P30. (D'-G') Magnification of areas within dotted lines in D-G. (H'-K', H"-K") Magnification of areas within dotted lines in H-K. Scale bars, 50 μm.

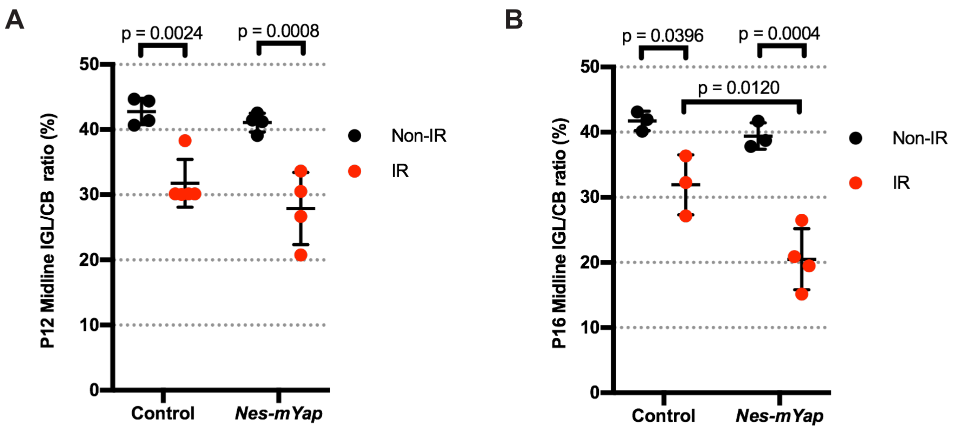


**Figure 2S2. The defect in recovery of mutants lacking *Yap* in NEPs from irradiation occurs after P12.**

(A-B) Graphs of the IGL/CB area ratios from *Nes-mYap* cKOs and controls at P12 (A) and P16 (B). P12 *Nes-mYap* cKO: Non-IR, n = 4; IR, n = 4; P12 control: n = 4; IR, n = 5; P16 *Nes-mYap* cKO: Non-IR, n = 3; IR, n = 3; P16 control: n = 3; IR, n = 3. Data are presented as mean ± S.D., and statistical analysis by two-way ANOVA. Each data point represents one animal.

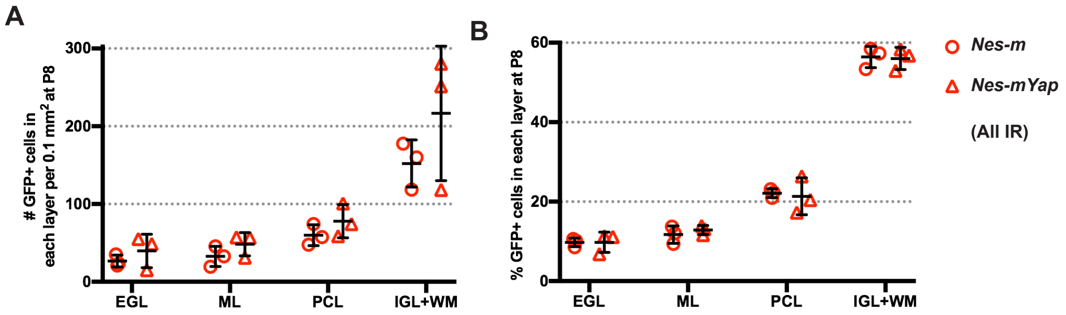


**Figure 3S. Deletion of of *Yap* in NEPs does not alter the distribution of Nestin-derived cells in the different layers of the cerebellar cortex at P8 after irradiation.**
Graphs of the numbers and percentages of GFP+ cells within each layer (A,B) and the total numbers of GFP+ cells (C) per 0.1 $mm^2$ of the total area analyzed in lobule 4/5 from midsagittal sections of IR *Nes-m* controls (n=3) and *Nes-mYap* cKOs (n = 3) at P8. Data are presented as mean ± S.D., and statistical analysis by two-way ANOVA. Each data point represents one animal.

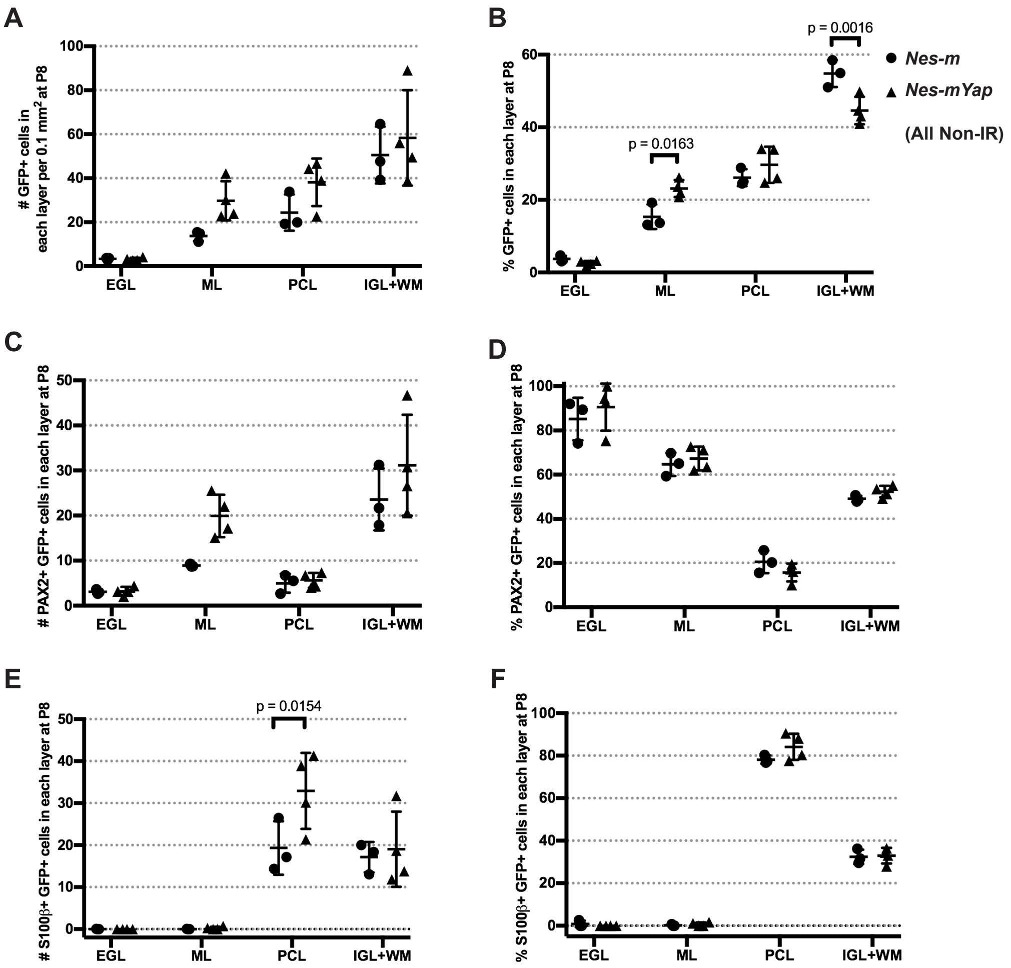


**Figure 4S1. YAP promotes differentiation of NEPs during normal postnatal CB development.**

Graphs of the numbers and percentages of GFP+ cells in the different layers (A,B), the numbers and percentages of PAX2+ GFP+ double cells (C,D) or S100β+ GFP+ double cells (E,F) within each layer, per 0.1 $mm^2$ of the total area analyzed in lobule 4/5 of midsagittal sections from *Nes-m* controls (n=3) and *Nes-mYap* cKOs (n = 4) at P8. Data are presented as mean ± S.D., and statistical analysis by two-way ANOVA. Each data point represents one animal.

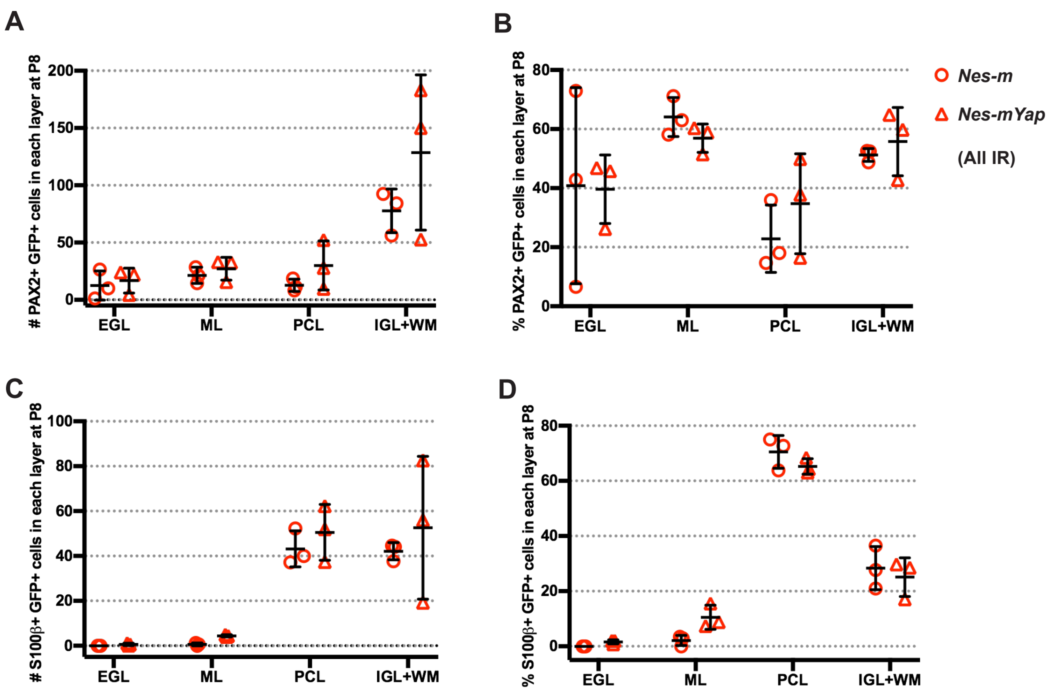


**Figure 4S2. Irradiation causing depletion of the EGL overrides the requirement for YAP in differentiation of NEPs.**

Graphs of the numbers and percentages of PAX2+ GFP+ double cells (A,B) or S100β+ GFP+ double cells (C,D) within each layer per 0.1 $mm^2$ of the total area analyzed in lobule 4/5 from midsagittal sections of IR *Nes-m* controls (n=3) and *Nes-mYap* cKOs (n = 3) at P8. Data are presented as mean ± S.D., and statistical analysis by two-way ANOVA. Each data point represents one animal.

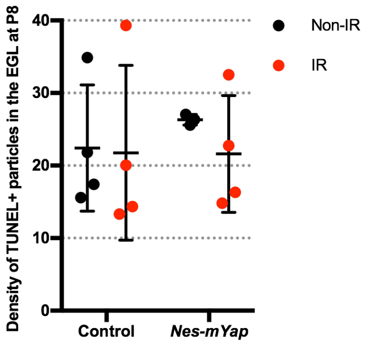


**Figure 5S. Loss of YAP does not increase cell death in the EGL at P8**
Graph of the densities of TUNEL+ particles in the EGL (the number of TUNEL+ particles per 0.1 $mm^2$ of EGL area) in midline CB sections from IR and Non-IR *Nes-mYap* cKOs and controls at P8. Controls: Non-IR, n = 4; IR, n = 4. *Nes-mYap* cKO: Non-IR, n = 3; IR, n = 4. Data are presented as mean ± S.D., and statistical analysis by two-way ANOVA. Each data point represents one animal.

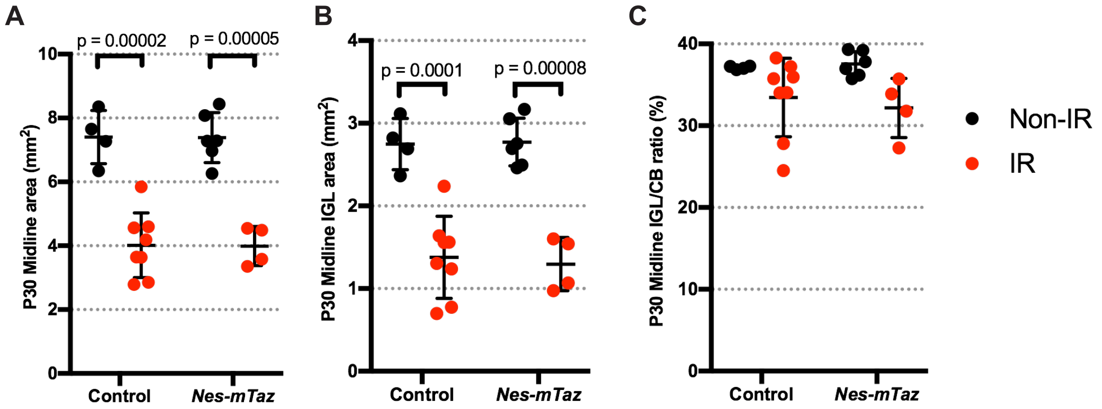


**Figure 7S. Loss of *Taz* in NEPs does not affect cerebellar growth during development and regeneration.**

(A-C) Graphs of the CB areas (A), the IGL areas (B), and IGL/CB area ratios (C) of midline CB sections from *Nes-mTaz* cKOs (Non-IR, n = 6; IR, n = 4) and controls (Non-IR, n = 4; IR, n = 8) at P30. Data are presented as mean ± S.D., and statistical analysis by two-way ANOVA. Each data point represents one animal.

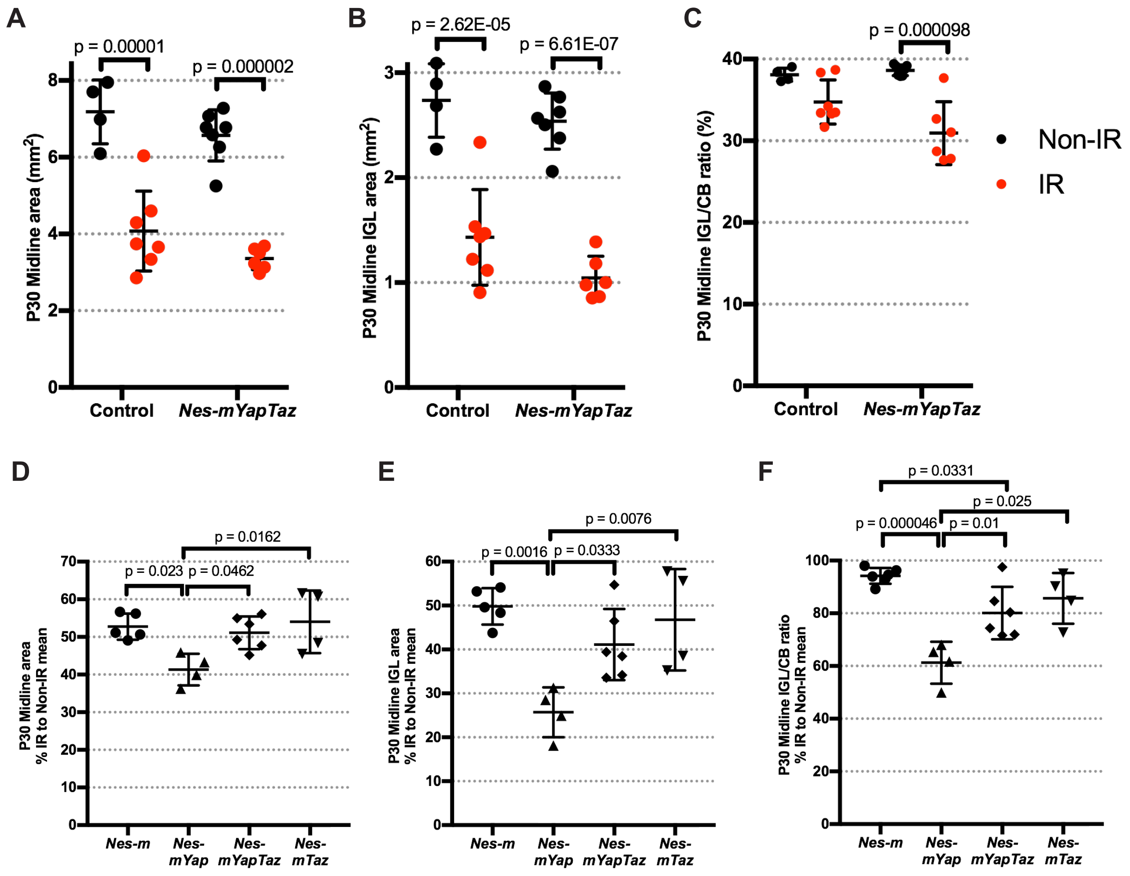


**Figure 8S1. Loss of *Taz* in NEPs lacking *Yap* does not abrogate recovery of the IGL after irradiation.**

(A-C) Graphs of the midline cerebellar areas (A), IGL areas (B), and IGL/CB ratios for IR and Non-IR *Nes-mYazTaz* cKOs (Non-IR, n = 7; IR, n = 6) and controls (Non-IR, n = 4; IR, n = 7) at P30. Data are presented as mean ± S.D., and statistical analysis by two-way ANOVA. Each data point represents one animal. (D-F) Graphs of the percentages of midline cerebellar areas (D), IGL areas (E), and IGL/CB ratios (F) for IR mice of the indicated genotypes as a percentage of Non-IR mice of the same genotype at P30. *Nes-m* controls (Non-IR, n = 6; IR, n = 5), *Nes-mYap* cKO (Non-IR, n = 6; IR, n = 4), *Nes-mYazTaz* cKO (Non-IR, n = 7; IR, n = 6), and *Nes-mTaz* cKO (Non-IR, n = 6; IR, n = 4). Data are presented as mean ± S.D., and statistical analysis by one-way ANOVA. Each data point represents one animal.

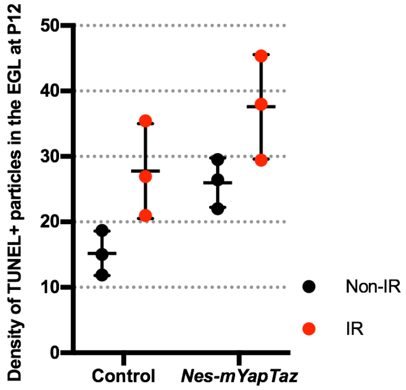


**Figure 8S2. Loss of *Yap* and *Taz* in NEPs at P0 does not result in a significant increase in cell death in the EGL at P12 after irradiation-induced EGL injury at P1.**
Graph of the densities of TUNEL+ particles in the EGL (the number of TUNEL+ particles per 0.1 $mm^2$ of EGL area) in midline cerebellar sections from IR and Non-IR *Nes-mYapTaz* cKOs and controls at P12. For all groups, n = 3. Data are presented as mean ± S.D., and statistical analysis by two-way ANOVA. Each data point represents one animal.
